# Effect of 10 UV filters on the brine shrimp *Artemia salina* and the marine microalgae *Tetraselmis* sp

**DOI:** 10.1101/2020.01.30.926451

**Authors:** Evane Thorel, Fanny Clergeaud, Lucie Jaugeon, Alice M. S. Rodrigues, Julie Lucas, Didier Stien, Philippe Lebaron

## Abstract

The presence of pharmaceutical and personal care products’ (PPCPs) residues in the aquatic environment is an emerging issue due to their uncontrolled release, through grey water, and accumulation in the environment that may affect living organisms, ecosystems and public health. The aim of this study is to assess the toxicity of benzophenone-3 (BP-3), bis-ethylhexyloxyphenol methoxyphenyl triazine (BEMT), butyl methoxydibenzoylmethane (BM), methylene bis-benzotriazolyl tetramethylbutylphenol (MBBT), 2-Ethylhexyl salicylate (ES), diethylaminohydroxybenzoyl hexyl benzoate (DHHB), diethylhexyl butamido triazone (DBT), ethylhexyl triazone (ET), homosalate (HS), and octocrylene (OC) to marine organisms from two major trophic levels including autotrophs (*Tetraselmis sp*.) and heterotrophs (*Artemia salina*). In general, EC_50_ results show that both HS and OC are the most toxic for our tested species, followed by a significant effect of BM on *Artemia salina* but only at high concentrations (1 mg/L) and then an effect of ES, BP3 and DHHB on the metabolic activity of the microalgae at 100 μg/L. BEMT, DBT, ET, MBBT had no effect on the tested organisms, even at high concentrations (2mg/L). OC toxicity represent a risk for those species since it is observed at concentrations only 15 to 90 times higher than the highest concentrations reported in the natural environment and HS toxicity is for the first time reported on microalgae and was very important on *Tetraselmis sp.* at concentrations close to the natural environment concentrations.

## Introduction

In recent decades, the sunscreen production and skin application have continuously increased to protect against sunlight exposure damages and to prevent skin cancer risk (Azoury and Lange, 2014; Waldman and Grant-Kels, 2019). Sunscreen products contain many chemical ingredients including UltraViolet (UV) filters, the aim of which is to absorb or reflect UVA and/or UVB radiations ranging from 280 to 400 nm (Sánchez-Quiles and Tovar-Sánchez, 2015).

More than 50 different UV filters are currently on the market (Shaath, 2010). Despite the fact that the use of these compounds is subject to different regulations around the world, UV filters are regularly detected in various aquatic environmental compartments including lakes, rivers, surface marine waters and sediments. These chemicals can enter in the marine environment in two ways, either indirectly from wastewater treatment plants effluent or directly from swimming or recreational activities (Giokas et al., 2007). Furthermore, their lipophilic nature results in low solubility in water, high stability and tendency to bioaccumulate (Gago-Ferrero et al., 2015; Vidal-Liñán et al., 2018).

In recent years, many ecotoxicological studies have focused on the impact of organic UV filters on various trophic levels, from microalgae to fish, passing by corals. Several studies have demonstrated that some of these compounds can disrupt survival (Chen et al., 2018; He et al., 2019; Paredes et al., 2014), behavior (Araújo et al., 2018; Barone et al., 2019), growth (Mao et al., 2017; Paredes et al., 2014; Sieratowicz et al., 2011), development (Giraldo et al., 2017; Torres et al., 2016), metabolism (Esperanza et al., 2019; Seoane et al., 2017; Stien et al., 2019), gene expression (Gao et al., 2013; Zucchi et al., 2011) and reproduction (Araújo et al., 2018; Coronado et al., 2008; Kaiser et al., 2012) in various species. It should be noted that the majority of toxicological studies on organic UV filters were conducted on BP3, EMC and 4-MBC.

The adoption and implementation of the European legislation on the registration, evaluation, authorization and restriction of chemicals (REACH) required several additional ecotoxicity data promoting the use of invertebrates as models for toxicity assays (European Commission, 2007). Brine shrimps *Artemia* spp. (here *A. salina*) are readily available worldwide and easy to breed. They have therefore been frequently used as test organism in ecotoxicity assays, and *A. salina* was chosen to investigate the toxicity of UV filters in this study (Caldwell et al., 2003; Libralato, 2014; Nunes et al., 2006).

The green algae *Dunaliella tertiolecta* is commonly used for chronic algal toxicity testing. The Haptophyta *Isochrysis galbana* is interesting too as it is widely cultured as food for the bivalve industry. Green algae such as *Chlorella* and *Tetraselmis* sp., belonging to the phyllum Chlorophyta, have also been frequently exploited in toxicity assays. *Tetraselmis* have been used before to study the toxic effect of several antibiotics and PCPs, but also the UV filter BP-3 (Seoane et al., 2017, 2014). This is why this algae was used in this study.

Previous studies about the toxic effects of different pollutants on microalgae physiology demonstrate that flow cytometry (FCM) can be an alternative to standard algal population-based endpoints, since it allows a rapid, quantitative and simultaneous measurement of multiple responses to stress in individual cells (Esperanza et al., 2019; Hadjoudja et al., 2009; Prado et al., 2015; Seoane et al., 2017). Seoane et al., (2017) showed that the most sensitive parameters are the metabolic activity and cytoplasmic membrane potential. Therefore, we decided to use FCM to analyze the toxicity of UV filters on *Tetraselmis* cells.

## 2. Materials and methods

### 2.1 Test substances and experimental solutions

The UV filters benzophenone-3 (BP-3), bis-ethylhexyloxyphenol methoxyphenyl triazine (BEMT), butyl methoxydibenzoylmethane (BM), and methylene bis-benzotriazolyl tetramethylbutylphenol (MBBT) were purchased from Sigma-Aldrich (Saint-Quentin Fallavier, France). 2-Ethylhexyl salicylate (ES), diethylaminohydroxybenzoyl hexyl benzoate (DHHB), diethylhexyl butamido triazone (DBT), ethylhexyl triazone (ET), homosalate (HS), and octocrylene (OC) were provided by Pierre Fabre Laboratories.

Before each toxicity test, and due to the low water solubility of the compounds, stock solutions at 1 mg/ml were prepared by dissolving each UV filters in dimethyl sulfoxide (DMSO, Sigma-Aldrich, purity >99%). These solutions were diluted in order to add the same amount of DMSO to all samples and to obtain exposure concentrations ranging from 20 ng/L to 2 mg/L for *A. salina*, and 10 μg/L to 1 mg/L for *Tetraselmis* sp.. The lower concentrations tested were roughly those reported in natural ecosystems. DMSO concentration in the experiments was always 2.5 % (v/v). A DMSO control (2.5 % v/v) and a blank control were also included. The blank control was artificial seawater for *A. salina* and growth medium for *Tetraselmis* sp..

### 2.2 Artemia salina mortality test

*A. salina* cysts were purchased from AquarHéak Aquaculture (Ars-en-Ré, France) and stored at 4 °C. Dried cysts were hatched in a constantly aerated transparent «V» hatching incubator filled with 500 mL of artificial seawater (ASW) at a salinity of 37 g/L, prepared with Instant Ocean salt (Aquarium Systems, Sarrebourg, France). Incubation was carried out for 48 h, at 25 °C under continuous light the first 24 hours. A 12:12 h light regime was then applied until the nauplii reached the instar II-III stage. Ten nauplii were transferred into 5 ml glass tubes filled with ASW (2 mL) contingently supplemented with DMSO and UV filters. The tubes were incubated at 25 °C under a 12:12 h light regime. The experiments were performed in sextuplicate. During the exposure period, there was no aeration and the nauplii were not fed. The mortality rate was estimated after 48 h by counting the dead nauplii under binocular. Organisms with no swimming activity or movement of appendices for 10 s even after mechanical stimulation with a Pasteur pipette were counted as dead. The tests were considered valid if the control’s average mortality rate was < 20%.

### 2.3 Tetraselmis sp. toxicity test

#### 2.3.1 Experimental procedure

*Tetraselmis* sp. (RCC500) was purchased from the Roscoff culture collection and was grown in filtered (pore size: 0.22 μm) and autoclaved seawater enriched with a 50-fold diluted f/2 medium (Sigma–Aldrich). The culture was maintained under controlled conditions at 18 °C (± 1 °C) with a photon flux of 70 μmol photons.m^−2^.s^−1^ under a dark:light cycle of 12:12 h. Toxicity tests were conducted in 150 mL Erlenmeyer flasks containing 50 mL of culture. Algae cells in exponential growth phase were used as inoculum and the initial cell density was 5.10^4^ cells/mL. Three replicates per UV filter concentration were performed. After 7 days of exposure, different morphological and physiological cells properties were monitored *via* flow cytometry (FCM). Analyzed parameters were granularity, relative cell volume, chlorophyll a fluorescence, esterase activity, and growth. The control experiment was a *Tetraselmis* sp. culture supplemented with DMSO (2.5%).

#### 2.3.2 Flow cytometry (FCM) analyses

Aliquots were collected after 7 days of exposition to be analyzed in a FACSCanto II flow cytometer (Becton Dickinson, Franklin Lakes, New Jersey, USA) equipped with an air-cooled argon laser (488 nm, 15 mW). To characterize the microalgae population, and to exclude non-algal particles, the forward scatter (FSC, an estimation of cell size) and side scatter (SSC, an estimation of granularity) dot-plots were used before each measurement. The flow rate of the cytometer was set to low (acquisition time: 1 min).

The data recorded by FCM were measured either directly (autofluorescence, granularity, size) or indirectly by the use of fluorochromes (esterase activity). Cellular density was determined using Becton Dickinson Trucount™ 10 μm beads for calibration, as described by Pecqueur et al. (2011). Growth rate (μ), expressed as day^-1^ were calculated using the following equation: μ =(ln(N_t_)−ln(N_0_)) / (t−t_0_), where N_t_ is the cell density at time t and N_0_ is initial cell density. Chlorophyll a natural autofluorescence was measured and detected in the FL3 channel (nm). Relative cell volume (or size) and granularity were directly estimated with the forward light scatter (FSC channel) and with the side scatter channel (SSC), respectively. To determine the metabolic activity based on the esterase activity study, cells were stained with the fluorochrome Chemchrom V6 (10-fold diluted in ChemSol B26 buffer– Biomérieux, France) at 1 % final concentration, and incubated for 15 min at room temperature in the dark before analysis. Reading was performed with the FL1 channel (nm). All cytometry data were analyzed using BD FACSDiva (Becton Dickinson). Results were expressed as percentage of variation relative to control (100 %).

### 2.4. Statistical analysis

Results are reported as mean and standard deviation (SD), calculated from the 3 or 6 replicates. For both tests and all the parameters measured, differences between controls and nominal concentrations of UV filter were analyzed using R software, by one-way analysis of variance (ANOVA) followed by post-hoc Tukey HSD tests for pairwise comparisons. In all cases, significance was accepted when *p* < 0.05. Dose–response curves, LD/LC_50_-values were estimated by a log(agonist) vs. response - Variable slope (four parameters) regression model in GraphPad Prism 5.

## 3. Results and discussion

### 3.1 Effects on Artemia salina mortality

The toxicity of several organic UV filters on the marine crustacean *Artemia salina* (Nauplii Instar II III) was determined after 48 h of exposure by counting dead larvae (Fig. 1). At the highest concentration tested (2 mg/L), HS, BM, and OC demonstrated a significant effect on Nauplii survival (p < 0.05) with mortality values reaching 54 ± 16%, 64 ± 19% and 88 ± 16%, respectively. At lower concentrations of these filters no significant effect was detected. For BP3, BEMT, MBBT, ES, DHHB, DBT and ET no toxicity was observed, even at the highest concentration.

**Fig 1:**
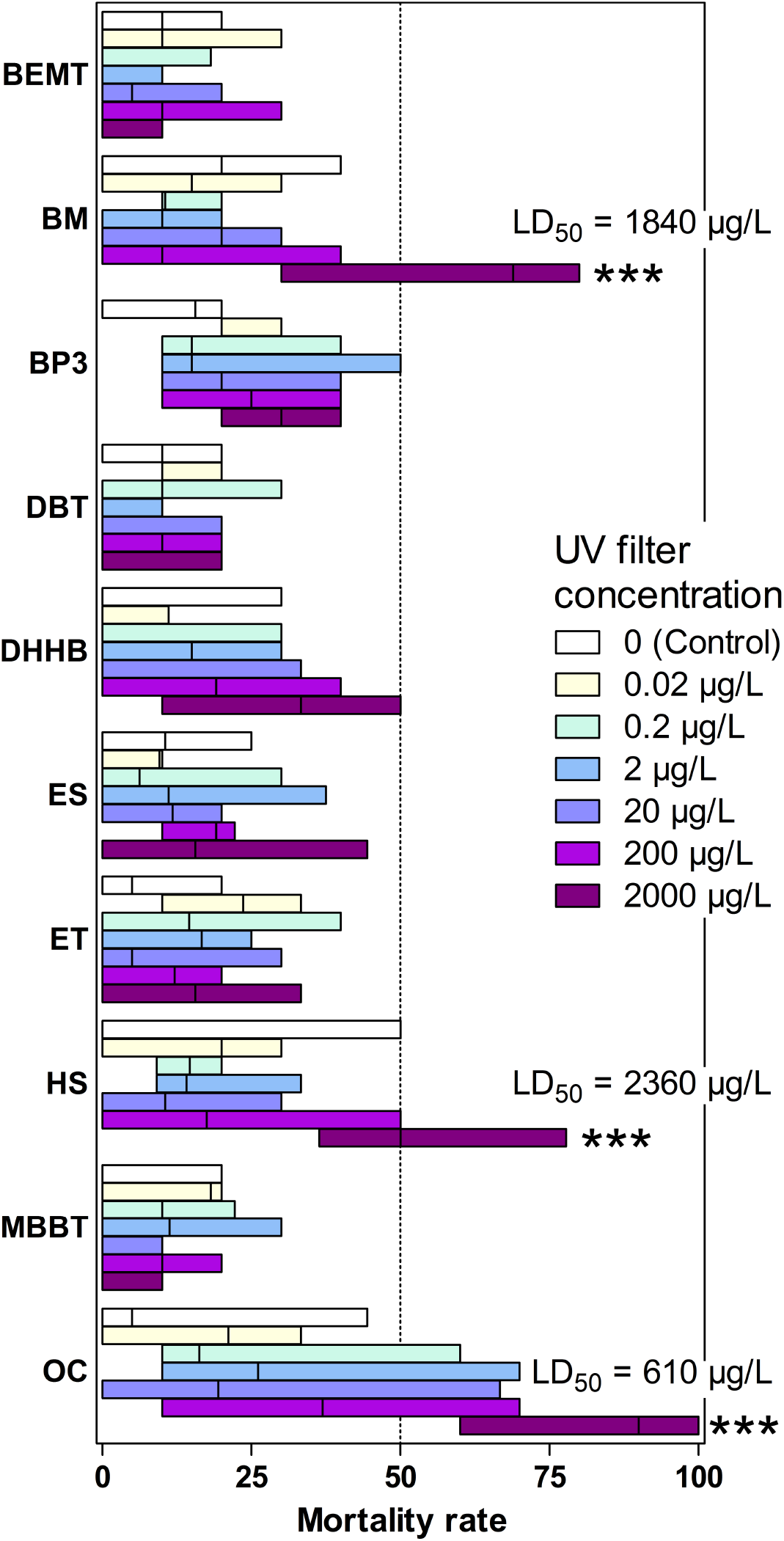
Mortality rate of *A. salina* exposed to the 10 UV filters at 6 concentrations. Boxes delineate the minimal and maximal values, and the vertical line is the median of six replicates. Significance levels relative to control determined by ANOVA followed by the Tukey’s multiple comparison test: *** p < 0.001. Results were not significant unless otherwise stated. For BM, HS and OC, the LD_50_ is reported on the figure.

Our results indicate that among the different UV filters tested in this study, OC was the most toxic molecule showing the lowest LD_50_ value (0.6 mg/L), followed by BM and HS (1.8 mg/L and 2.4 mg/L, respectively). Environmental HS and BM concentrations reported so far are at least 500 times lower than LD_50_, with values always lower than 3 μg/L (Fagervold et al., 2019; Sánchez-Quiles and Tovar-Sánchez, 2015; Tsui et al., 2014; Ramos et al., 2015). OC concentrations in coastal waters are higher and can reach 9 μg/L (Langford and Thomas, 2008; Tsui et al., 2014; Ramos et al, 2015). This is the first report showing OC toxicity on *Artemia salina*. We also observed a concentration-dependent increase in mortality of *Artemia* with respect to the control. This is congruent with the toxicity observed, at lower concentration, on coral (50 μg/L, (Stien et al., 2019), urchin, mussel and algae (40-80 μg/L, (Giraldo et al., 2017). OC also affects the developmental process in zebrafish (Blüthgen et al., 2014). Here the LD_50_ on *A. salina* is approximately 90 times higher than the highest OC concentrations in marine waters reported so far. It should be mentioned as well that OC concentrations in the 50-100 μg/kg range have been frequently reported in sediments (Gago-Ferrero et al., 2011; Kameda et al., 2011), in which case OC may affect benthic crustaceans.

### 3.2 Effects on Tetraselmis sp

#### 3.2.1 Growth rate and EC_50_ values

After 7 days of exposure, HS, BP3, and ES induced a significant decrease of algae growth (Fig. 2A). The growth rate of algae exposed to ES at 1 mg/L decreased by 24 % compared to control (p < 0.05). For BP-3 we observed a concentration dependent decrease in growth, which was statistically significant at 100 μg/L (p < 0.01) and 1 mg/L (p < 0.001). At 1 mg/L, the growth rate was negative, which translates a decreased cell concentration compared to t_0_. The 7-days LC_50_ value for BP3 was 143 μg/L. BP3 LC_50_ values on several microalgae species have been reported previously to be roughly in the 100 μg/L to 20 mg/L range (Esperanza et al., 2019; Mao et al., 2017; Pablos et al., 2015; Zhong et al., 2019). Seawater BP3 concentrations in the μg/L range have been frequently reported in the literature (Gago-Ferrero et al., 2015; Tsui et al., 2014). Extremely high values of 1.4 and 0.6 mg/L have been recorded in the U.S. Virgin Islands (Downs et al., 2016). Finally, the most important decline was with UV filter HS. No algal cells were detected in the presence of HS at 100 μg/L and 1 mg/L. LC_50_ with HS was estimated at 74 μg/L, while it has been shown that HS concentration in aquatic environments can reach ~3 μg/L (Tsui et al. 2014; Ramos et al., 2015). OC induced a slight but significant increase of the growth rate at 1mg/L with a decrease in the metabolic activity as determined by esterase activity. This increased metabolic activity was not explained. Similar differences in the response of different physiological parameters were already reported by Esperanza et al (2019) for the toxicity of BP3 in the microalgae *Chlamydomonas reinhardtii*. The growth rate of *Tetraselmis* was not affected with BEMT, MBBT, DHHB, DBT, ET and BM, even at 1 mg/L.

**Fig 2:**
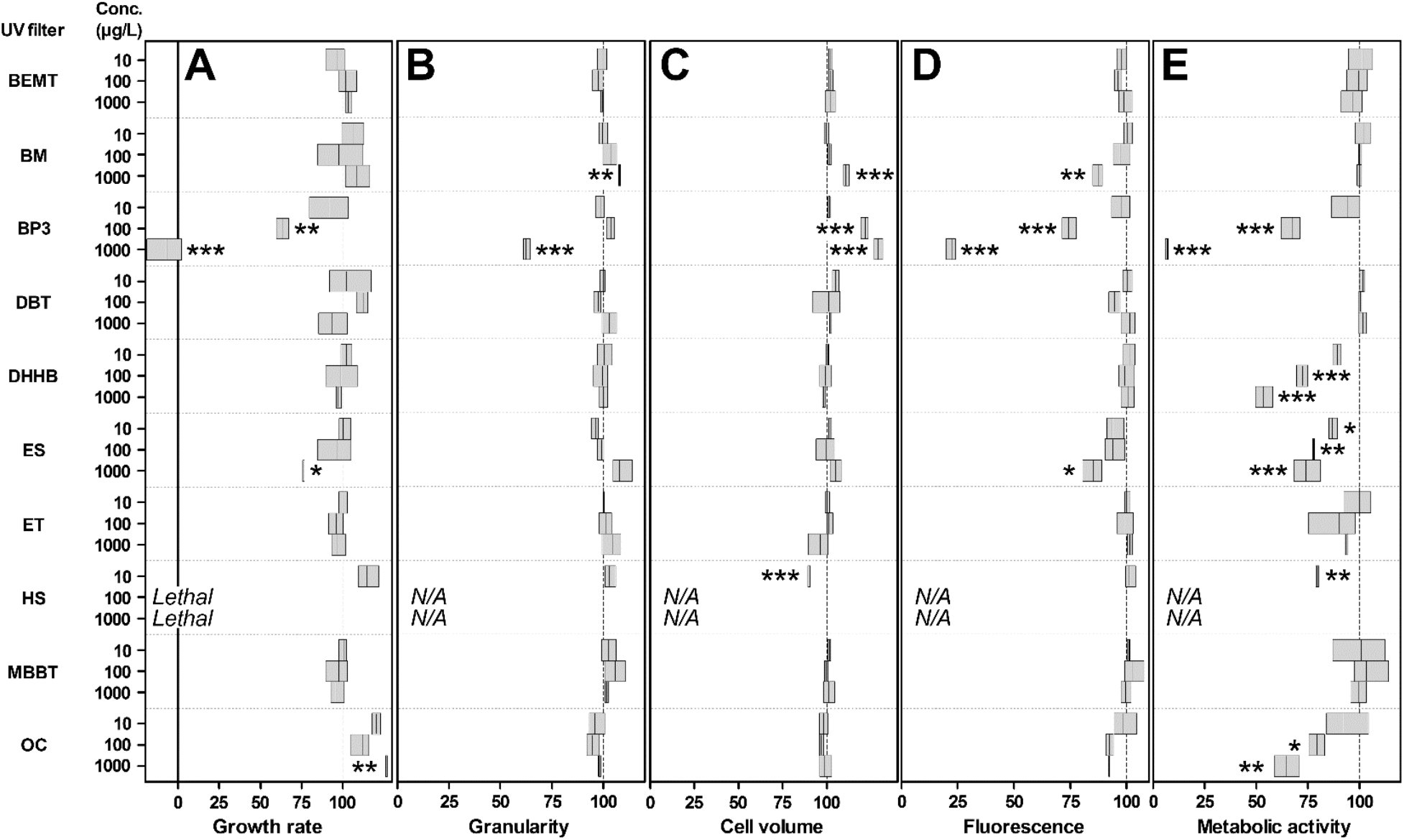
Relative A) growth rate, B) granularity, C) cell volume, D) fluorescence and E) metabolic activity of exposed *Tetraselmis* compared to control, set to 100 %. The boxes delineate the minimal and maximal values. The vertical line in the boxes is at mean. Significance levels relative to negative control determined by ANOVA followed by the Tukey’s multiple comparison test: *** p < 0.001, ** p < 0.01, * p < 0.05. Results were not significant unless otherwise stated. N/A: not applicable, the data could not be obtained due to extensive cell death.

#### 3.2.2 Impact on cell morphology

Three of the UV filters induced cell morphological alterations (Fig. 2B/C). Cells cultured in the presence of BM have experienced a significant increase in cell volume and granularity at 1 mg/L (p < 0.05). This concentration is 1,000 to 10,000 times higher than the few concentrations reported in the field (Fagervold et al., 2019; Sánchez-Quiles and Tovar-Sánchez, 2015; Tsui et al., 2014). According to environmental concentrations of other UV filters, one can assume that the effective concentration of 1 mg/L is probably higher than any BM environmental concentration, but this remains to be confirmed. BP3 caused a dose dependent increase of relative cell volume at 100 μg/L and above, reaching up to 129 % of control cell volume at 1 mg/L (p < 0.001). Meanwhile, this UV filter induced a significant 38 % granularity decrease at 1 mg/L (p < 0.001). With reference to the environmental concentrations of BP3 (see above), this filter should exert a significant impact on phytoplankton communities. These results are congruent with what was recently reported by Zhong et al. (2019) on *Arthrospira* sp. With HS, cell volume and granularity could not be measured at 100 and 1000 μg/L. However, a significant cell volume decrease was observed at 10 μg/L of HS (−11 %, p < 0.001). In our experiment, HS No Observed Effect Concentration (NOEC) was therefore lower than 10 μg/L, *i.e.*, within the same order of magnitude than the highest water column concentration reported so far by Tsui et al. (2014). Again, it is expected that HS should affect microalgae communities in bathing areas. No significant effect was observed for BEMT, MBBT, ES, DHHB, DBT, ET, HS and OC.

#### 3.2.3 Impact on autofluorescence

The results of FCM analysis revealed that several UV filters significantly reduced chlorophyll a (Chl a) cell fluorescence (Fig 4D). The decrease was significant with BM (–13 %, p < 0.01) and ES (−15 %, p < 0.05) at 1 mg/L. A strong dose-dependent autofluorescence inhibition was observed upon exposure to BP3 at concentrations of 100 μg/L (p < 0.001) and above. Inhibition reached 78 % at the highest dose. Again, autofluorescence could not be measured in cells treated with HS at 100 and 1000 μg/L due to the important cell degradation at these concentrations. No significant effect was observed for BEMT, MBBT, DHHB, DBT, ET and OC.

#### 3.2.4 Impact on cell metabolic activity

Metabolic activity was determined by estimating the relative esterase activity in exposed cells compared to control ones. It was measured by CV6 staining and highlighted significant decreased metabolic activities with half of the tested UV filters (Fig 2E). Algae exposed to ES and HS experienced a decreased esterase activity at 10 μg/L UV filter. The effect of BP3, DHHB and OC was significant at 100 μg/L and above. A similar decreased esterase activity was reported for *C. reinhardtii* exposed to BP3, although at concentrations in the mg/L range (Esperanza et al., 2019; Seoane et al., 2017). For DHHB, the effect was only observed for the microalgae and for the esterase activity but not for other parameters. Therefore, the environmental risk cannot be estimated since natural concentrations have never been reported for this filter. No significant effect was observed for BEMT, MBBT, DBT, ET and BM.

## 4. Conclusion

The present work demonstrates that several filters exert toxicity on *A. salina* and *Tetraselmis* sp. HS was the most toxic UV filter for the microalgae. LC_50_ was 74 μg/L and significant adverse effects were recorded at the lowest concentration tested (10 μg/L). HS was also toxic for *A. salina*, although at much higher concentrations (LD_50_ 2.4 mg/L). Overall, HS NOEC was lower than 10 μg/L. Its Lowest Observed Effect Concentration (LOEC) was 10 μg/L or below. HS concentrations up to 3 μg/L have been reported in the natural environment in which case HS may represent a potential risk for marine phytoplankton communities. Further research is needed to investigate on HS toxicity with a larger diversity of phytoplankton species.

OC was toxic on both models with a dose-dependent effect on the microalgae. OC significantly altered *Tetraselmis* sp. metabolic activity at 100 μg/L. On *A. salina*, LD_50_ was 610 μg/L. Overall, OC LOEC was 100 μg/L with these models. The toxicity of OC occurred at concentrations only 90 (*A. salina*) and 15 (*Tetraselmis* sp.) times higher than the highest environmental concentrations reported so far. These results confirm the toxicity of OC on marine organisms. BM was toxic towards the brine shrimp at high concentrations with a LD_50_ of 1.84 mg/L and had little effect on the microalgae. BM LOEC was 1 mg/L. The toxic concentrations reported here may be high enough. BM might not have any effect on marine ecosystems, although the occurrence of this filter should be monitored in a large range of ecosystems to better estimate its natural concentrations. ES (LOEC 10 μg/L), BP3 and DHHB (LOEC 100 μg/L) had a significant impact on the microalgae metabolic activity and had little effect on *A. salina*.

Overall, this research supports the need of establishing environmental quality standards for UV-filters based on toxicity testing with key marine organisms, as well as identifying and reducing environment input sources for the toxic ones. There are still many filters for which environmental concentrations are missing to better estimate the potential environmental risk of their occurrence in coastal ecosytems. It is probably important also to design new user’s and environment friendly UV filters.

## Acknowledgements

We thank the BIO2MAR and the BIOPIC platforms from the Observatoire Océanologique de Banyuls for providing technical support and access to instrumentation.

